# Powered flight potential approached by wide range of close avian relatives but achieved selectively

**DOI:** 10.1101/2020.04.17.046169

**Authors:** R. Pei, M. Pittman, P.A. Goloboff, T.A. Dececchi, M.B. Habib, T.G. Kaye, H.C.E. Larsson, M.A. Norell, S.L. Brusatte, X. Xu

**Author notes:** These authors contributed equally to this paper (R. Pei, M. Pittman). Correspondence and requests for materials should be addressed to M.P.

## Abstract

Evolution of birds from non-flying theropod dinosaurs is a classic evolutionary transition, but a deeper understanding of early flight has been frustrated by disagreement on the relationships between birds (Avialae) and their closest theropod relatives. We address this through a larger, more resolved evolutionary hypothesis produced by a novel automated analysis pipeline tailored for large morphological datasets. We corroborate the grouping of dromaeosaurids + troodontids (Deinonychosauria) as the sister taxon to birds (Paraves), as well as the recovery of Anchiornithidae as basalmost avialans. Using these phylogenetic results and available data for vaned feathered paravians, maximum and minimum estimates of wing loading and specific lift calculated using ancestral state reconstruction analysis are used as proxies for the potential for powered flight through this transition. We found a broad range of paravian ancestors with estimates approaching values that are indicative of powered flight potential. This suggests that prior to the evolution of flight there was a wider extent of experimentation with wing-assisted locomotion among paravians than previously appreciated. We recovered wing loading and specific lift estimates indicating the potential for powered flight among fossil birds as well as unenlagiine and microraptorine dromaeosaurids. In the context of our phylogeny and of Mesozoic palaeogeography, our results suggest that the potential for powered flight originated three or more times from a broad range of ancestors already nearing this potential, providing a well-supported scenario for the origin of theropod flight to further explore.

---

The origin of birds (Avialae) and modern powered flapping flight were iconic events in the history of life. Recent studies of early birds and their closest dinosaurian relatives (non-avialan paravian theropods) have provided key insights into this major evolutionary transition. It is now clear that anatomies and behaviours traditionally associated with birds were first acquired by non-avialan dinosaurs before the origin of birds and modern powered flapping flight. These include smaller body size, accelerated evolutionary rates^1-3^, early feathers of ‘modern’ aspect^4-17^, complex plumage colouration, flapping-based locomotion, non-powered flight capabilities among some non-avialan paravians^18-24^, and even an avian-like sleeping posture^25^. As a result of these advances, the origin of birds has emerged as one of the best documented examples of a major macroevolutionary transition. Despite the extensive array of new specimens and data, the phylogenetic relationships within and between avialans, dromaeosaurids and troodontids have remained unclear with both a lack of consensus and resolution at important nodes.

Traditionally, dromaeosaurids and troodontids were united together as the Deinonychosauria by the ‘sickle-claw’ of their second toe and other characters^2,26-29^. They were considered the sister group of birds and were altogether known as the Paraves^2,26-29^. The rapid discovery of paravian species over the last decade^13,14,17,29-33^, especially from East Asia, has called this into question. Most notably, many of the historically diagnostic features of Dromaeosauridae, Troodontidae and Deinonychosauria are now recognised as synapomorphies of more inclusive theropod groups (e.g. Maniraptora), or in some cases appear to have been acquired convergently in different taxa. The number of evolutionary hypotheses has grown with these new fossil discoveries^1,2,13,14,29,31,33-39^. They now encompass almost every possible combination of interrelationships between birds and other paravians, even challenging the monophyly of Deinonychosauria and the composition of stem avialans^13,34-36,39,40^. A primary issue concerns troodontids which have been grouped with either dromaeosaurids^2,29,33,34,39^ as the traditional Deinonychosauria, or with Avialae^13,14,35,36^ exclusive of dromaeosaurids. Each phylogenetic hypothesis has different implications on the origin of birds and the morphological, biomechanical and ecological states of their transitional antecedents.

We address these phylogenetic issues by presenting an updated parsimony-based reconstruction of paravian interrelationships using a large, long-standing coelurosaur theropod dataset (^1^ and references therein) expanded with recently discovered taxa. This dataset includes nine new dromaeosaurid terminal taxa (*Acheroraptor, Changyuraptor, Dakotaraptor*, IVPP V22530, *Linheraptor, Luanchuanraptor, Velociraptor osmolskae, Yurgovuchia* and *Zhenyuanlong*) giving the largest total number of dromaeosaurids (31) used so far in a phylogenetic analysis. It also incorporates a wealth of new data from existing paravians that have been recently described in more detail^15-17,29,31-33,36,41^ and studied first-hand, including the key basal paravians *Anchiornis* and *Archaeopteryx*^42^. For additional details see ‘Phylogenetic dataset’ in Methods.

Palaeontological datasets often pose important challenges to phylogenetic analysis, especially in terms of taxa that can be placed equally well in distant parts of the tree (“wildcards”), typically as a consequence of missing entries from incomplete preservation. Further challenges may emerge from high degrees of morphological variation, manifesting as homoplasies, in densely sampled phylogenetic regions. Both of these confounding issues are expected to be present in bird origin studies. A key goal of this study is to provide accurate phylogenetic placement for as many paravians as possible, but several are missing over 90% of their scoring entries (e.g. the dromaeosaurids *Atrociraptor*, and *Shanag*). We have thus placed special emphasis in developing a pipeline of analysis that can automatically deal with the numerous wildcards commonly seen in palaeontological datasets, including some newly developed techniques. The steps in a phylogenetic analysis subsequent to finding the optimal trees, particularly when wildcards are involved, are prone to human error (which, with many of the steps being sequential, may easily carry over to the following steps). Thus, to minimise the risk of human error, scripts have been used to automate all analytical procedures, in a way that is both reproducible and appropriate for other palaeontological datasets (with only minor modifications). Perhaps the main benefit of automated analysis is it encourages the dataset to be rechecked and corrected for scoring errors or problems. This analytical pipeline is therefore an important product of this study with its enhanced automation and newly developed techniques, and should greatly increase access to more in-depth analyses using parsimony as a criterion for phylogenetic inference.

We use this well-resolved phylogeny to infer when and how flapping-based locomotion evolved in the closest relatives of birds, specifically when and how the potential for powered flight developed. Previous work has proposed that powered flight in theropods evolved once or maybe even multiple times. It has even been suggested that birds should be defined by the possession of flight alone, as an apomorphic feature^43^. With the phylogenetic placement of the iconic early bird *Archaeopteryx* with Deinonychosauria, a single-origin of powered flight has been proposed at Paraves, polarizing the evolution of proportionally longer and more robust arms at that node^39^. However, the wing and body dimensions of many basal paravians do not surpass the minimal thresholds for flight ability as defined in modern birds and other taxa^44-46^. A previous quantitative study found that non-volant flapping-based locomotion was confined to Paraves: flap running, wing assisted incline running [WAIR], and wing-assisted leaping^18^. In showing that this was optimised at Paraves and that significant capabilities were derived independently in microraptorine dromaeosaurids and avialans, that study supported the potential for multiple origins of powered flight in theropods^18^. However, that study was restricted by a problematic phylogeny and small taxon sample, but more importantly, it did not focus on testing taxa and lineages against known minimal thresholds for flight ability in modern birds^44-46^.

We have overcome these restrictions by using the new, larger and more resolved tree topology across 43 taxa sharing lift-compatible vaned feathers (Pennaraptora; but see ^47^) to provide maximum and minimum estimates of wing loading and specific lift in the ancestors of our study taxa using ancestral state reconstruction analysis. These provide a proxy of the potential for powered flight through this transition from non-flying to flying theropods. These parameters are estimated from morphological features measurable from the fossils and are commonly used to evaluate flight capability in extant avians^44-46^. Wing loading is a major determinant of minimum flight speed, required launch and landing speeds, and manoeuvrability in powered flyers^48^. Wing loading is also a major determinant of required flapping frequency in powered flyers (wings must be flapped faster if they are smaller and/or body size is greater)^49^. In powered flyer, specific lift is critical to weight support and generation of thrust (thrust is primarily a component of lift in vertebrate flapping flyers)^50^. Whilst small powered flyers with sophisticated wing kinematics (particularly during the upstroke phase) can generate some drag-based thrust, we consider this to have been unlikely in basal birds because of their less refined aerofoils and motion control. The three major types of biomechanical competency for flight are assessed by wing loading and specific lift:

1. *Anatomical requirements for flight* Takeoff in flying animals is initiated by leaping^51^ so the primary anatomical requirements to initiate launch in paravians are related to hind limb characteristics^18^. Because of the cursorial ancestry of theropod dinosaurs, large hind limb muscle mass and robust hind limb skeletal elements were plesiomorphic for paravians, so all of the taxa we examined inherited sufficient hind limb strength for leaping^18^. For powered flapping flight (i.e. after takeoff), the primary anatomical requirements are summarized by wing loading, which simultaneously includes potential lift-producing surface area and body weight in one single variable. A key assumption we make is that our fossil taxa had body densities within the range known for living birds. We constrained the estimated body mass range of taxa potentially capable of powered flight by assuming that they were roughly similar in mass to living birds with similar wingspans and body volumes. This gave us a narrower set of body mass estimates within the relative large confidence intervals around the regressions used to estimate body mass^52^.
2. *Aerodynamic force production requirements for flight* Early in theropod flight evolution, the lift:drag ratios of wings were not necessarily equivalent to those of modern birds^20,53,54^. However, morphospace comparisons of wing shape show significant overlap between early taxa and modern ones^55^. There is no evidence that the airfoil shapes of fossil taxa differed significantly from living taxa e.g. fossil taxa also possessed similar leading edge shapes with well-developed propatagia^56-58^. The long bone cross sections in the forelimbs of early birds and microraptorine dromaeosaurids have similar shapes and comparable bending strengths to those of living birds^59^. Analysis of feather stiffness^60^ and vane asymmetry ratios^61^ demonstrate that the feathers of early paravians may have been less competent as individual airfoils than the primary feathers of living birds (but see ^62^). This may have limited earlier taxa to the use of unslotted wings. Furthermore, some questions remain regarding the upstroke kinematics available to early paravians^18,63,64^. Taken together, these data indicate that early paravians were capable of similar aerodynamic force production to that seen during steady-state conditions in living birds, excluding the use of slotted wing tips. In quantitative terms, these data suggest that lift coefficients up to 1.6 (typical steady state maximum for living birds) were possible, but the larger lift coefficients sometimes achieved by living taxa using dynamic stall and similar unsteady mechanisms (as high as 5.3 – see Norberg^65^) may not have been possible. Wing loading and specific lift estimates of fossil theropods that pass value thresholds characterizing all volant modern birds therefore indicate a potential for powered flight.
3. *Physiological requirements for flight* Our estimates of specific lift utilize a range of potential muscle power available to our theropod taxa to reflect the prevailing uncertainty in this parameter. We assume that at least some anaerobic power was available for climb out after takeoff, and we have included this in our estimates, but we have kept the estimates of this anaerobic fraction conservative (see Supplemental Information). The specific lift estimates also take into account the likely limitations on the maximum coefficient of lift in early taxa mentioned above.

Estimates of wing loading and specific lift were calculated from reconstructed ancestral morphologies using our own direct measurements of specimens as well as parameters reported in the literature. This allowed us to identify ancestors that fall within the range seen in extant volant birds, which we consider more accurate than just mapping measurements of flight capability. To consider parametric differences in past studies and differences from on-going uncertainties in paravian anatomy (see Methods), we calculated a maximum and minimum estimate for wing loading and specific lift. These estimates bracket the range of calculation permutations currently available, producing the most conservative results currently possible (for additional information see Methods and Supplementary Figs. S7-S10). Our approach contrasts with the concept of body weight support examined in ^18^, as we wanted to avoid using poorly known behavioural capabilities (e.g. flapping speed, flap angle and running speed) in our calculations in order to maximise our model precision. We interpret our results in the context of updated osteological and feather anatomy changes recovered from the revised phylogeny to provide the most detailed account yet of when and how powered flight evolved as a modification of flapping-based locomotion in theropod dinosaurs.

## Results & Discussion

### Paravian phylogeny

Our newly recovered phylogenetic details allow us to discuss paravian evolution in greater detail than previous studies. All of our extended implied weighting (XIW) and equal weighting (EW) topologies support the monophyly of each of the traditionally recognised paravian clades: Paraves comprises of Deinonychosauria and Avialae as in ^2,26-29^ and Deinonychosauria comprises of Dromaeosauridae and Troodontidae as in ^2,29,33,34,39^ (strict consensus and pruned reduced consensus trees using XIW [Figs. 1, S1, S3] and EW [Figs. 1, S2, S4]; see ‘Character weighting’ in Methods). Dromaeosaurid interrelationships are significantly improved relative to previous studies with well-supported internal resolution. The results gather the four anchiornithid taxa, which were previously scattered throughout Paraves^1,2^, into a distinct clade of basal avialans (Anchiornithidae) as in ^35,42,66^ and, in part ^39^ (Figs. 1, S1-S4). Our results allow us to present a revised list of evolutionary synapomorphies for the major paravian clades, including a refined sequence of evolutionary changes at the base of Avialae (Fig. 1). A succinct description of the results is given here (for additional details see Supplementary Note 1).

**Fig. 1.**
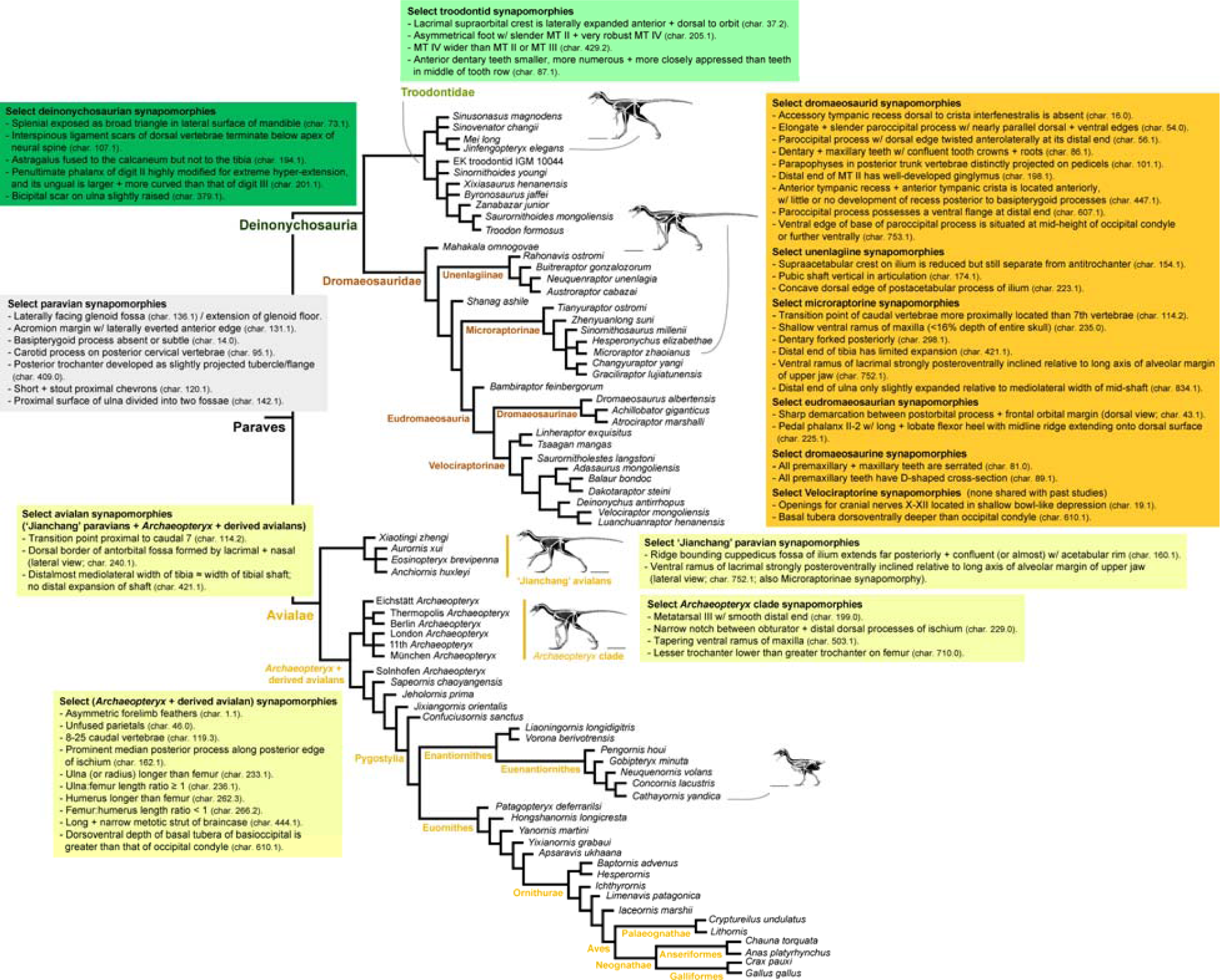
Revised paravian phylogeny and select nodal synapomorphies. Reduced strict consensus tree showing topology and select synapomorphies common to analyses using extended implied character weighting and equal character weighting. The synapomorphies shown are shared with past studies (see Results for additional information). For previously unreported synapomorphies see Supplementary Note 1. See Methods for additional information. Skeletal reconstructions (scale bar = 10cm) are used with the permission of Scott A. Hartman.

Despite differences with paravian interrelationships recovered in previous studies^1,2,13,14,29,31,33-39^, some of these past studies share the same synapomorphies for Paraves, making them useful traits for identifying members of the clade. A laterally facing glenoid fossa appears to be an especially useful trait for identification (character [char.] 136.1 in this study) as it was recovered at the equivalent node by Turner *et al*. ^2^ and Brusatte *et al.* ^1^ (char. 138.1 in ^2^ and char. 136.1 in ^1^) and is related to the extension of the glenoid floor onto the external surface of the scapula, which is a paravian synapomorphy of Agnolin & Novas^66^ (char. 138.1 in ^66^), Senter *et al*. ^29^ (char. 216.1 in ^29^) and Xu *et al*. (char. 122.1 in ^34^) as well as a synapomorphy of Foth *et al*. ^35^ at the node (*Jinfengopteryx* + Paraves) (char. 133.1 in ^35^).

The basic dromaeosaurid topology of previous analyses is maintained in this study with unenlagiines and microraptorines at more basal positions (*contra* ^66^) and eudromaeosaurians at more derived ones^1,2,15,17,29,31,33^. Regardless of the character weighting employed, *Mahakala* is recovered as the basalmost dromaeosaurid followed by the Unenlagiinae (*Rahonavis* sister to *Buitreraptor* and (*Neuquenraptor* + *Austroraptor*)) and then *Shanag*, with the latter taxon sister to the clade (Microraptorinae + Eudromaeosauria). Recent phylogenetic analyses have had limited success in resolving microraptorine interrelationships with Brusatte *et al*. ^1^, Turner *et al*. ^2^ and Lü & Brusatte^17^ all recovering polytomies in their reduced strict consensus trees (Fig. 64 of ^2^; Fig. S2 of ^1^; Fig. 5 of ^17^), whereas the topologies recovered by Senter *et al.* ^29^ and Xu *et al*. ^34^ were resolved. Under both XIW and EW results, we recover the largest microraptorine, *Tianyuraptor*, as the basalmost member of Microraptorinae, as suggested by Senter *et al*. ^29^, but contrary to its previously suggested eudromaeosaurian affinity^33^. We also recover another relatively large-bodied microraptorine, *Zhenyuanlong*, as the next most basal microraptorine for the first time, which was previously placed in a polytomy of Liaoning dromaeosaurids^17^). *Tianyuraptor* and *Zhenyuanlong* lack both a characteristic tubercle along the lateral edge of the mid-shaft of the posteriorly curved pubis and a subarctometatarsalian foot, unlike other microraptorines (chars. 228.0 and 200.1). However, *Tianyuraptor* is unique among microraptorines in having a straight pubis (char. 177.0), resembling other non-unenlagiine and non-velociraptorine dromaeosaurids. Other smaller derived microraptorines (char. 200.1) are recovered as a polytomy in both the XIW and EW strict consensus trees. As expected, the derived microraptorines include the Early Cretaceous Chinese dromaeosaurids *Changyuraptor* and IVPP V22530^15,32^.

*Bambiraptor* was recovered as the basalmost eudromaeosaurian in all XIW results, whereas in all EW results *Bambiraptor* is the second most basal eudromaeosaurian after *Saurornitholestes.* A relatively basal position for *Bambiraptor* within Eudromaeosauria was recovered by Agnolín & Novas^66^, Senter *et al.* ^29^ and DePalma *et al*. ^33^, whilst both *Bambiraptor* and *Saurornitholestes* were nested within Velociraptorinae according to Turner *et al*. ^2^ and Brusatte *et al*. ^1^. Under both equal and differential weighting, the remaining traditionally identified eudromaeosaurians do not have resolved interrelationships in the strict consensus tree except for *Linheraptor* and *Tsaagan*, which are recovered in a sister relationship as expected^41^. Pruning *Yurgovuchia, Acheroraptor, V. osmolskae* and *Utahraptor* from the eudromaeosaurian polytomy in both the XIW and EW strict consensus tree reveals a much more resolved topology, with a monophyletic Dromaeosaurinae and Velociraptorinae (See Supplementary Note 1).

Our results under XIW and EW fail to recover Anchiornithidae as part of Troodontidae (*contra* ^1,2^). The remaining troodontids were recovered in two clades as in the reduced strict consensus trees of Brusatte *et al.* ^1^ (Fig. S2 of ^1^: at least two clades) and in the strict consensus tree of Turner *et al.* ^2^ (Fig. 57 of ^2^), but their composition differs (see Supplementary Note 1).

In the XIW and EW topologies, the anchiornithid paravians from northeastern China – *Anchiornis, Aurornis, Eosinopteryx* and *Xiaotingia* – were recovered as the basalmost avialan clade (more basal than *Archaeopteryx* and derived avialans) as in ^35,42,66^ and, in part, ^39^. The avialan node recovered in this study is shared by the anchiornithids and traditional avialans.

Group support calculated by means of symmetric resampling^67^ (with character weights doubled or halved, never eliminated) were generally very low within Paraves, indicating that nodes are supported with relatively high levels of character conflict (Figs. S5,S6). Disregarding the most unstable taxa, reasonable support values between 76 and 100 were found for the remaining tree, including Paraves, Troodontidae, Dromaeosauridae, Unenlagiinae and Eudromaeosauria (Figs. S5,S6). In other datasets^1,2,29,34,35^, the nodal supports of paravian nodes are generally low, with Bremer supports typically between 1 and 2 for most paravian nodes (but see ^34^). See ‘Group supports and conflict’ in Supplementary Methods for more information.

### Major types of biomechanical competency for flight

Our new, better resolved phylogeny provides a firmer basis to assess the three major types of biomechanical competency for flight among paravian dinosaurs. To what extent do they meet the anatomical, aerodynamic force production and physiological requirements for flight we see in modern birds?

1. *Anatomical requirements for flight* The primary anatomical requirements for flapping flight are summarized by wing loading. All of the non-paravian vane-feathered theropods sampled have estimated wing loading values well above 2.5 gcm^−2^, with the lowest values of ∼6.0 gcm^−2^ (Figs. 2, S7, S8) being larger than values previously measured in extant flightless birds such as flightless ducks, cormorants, or Kakapos^68-70^. Using ancestral morphologies calculated using ancestral state reconstruction analysis, all avialans (including anchiornithids), six dromaeosaurids (*Bambiraptor, Buitreraptor, Changyuraptor, Mahakala, Microraptor* and *Rahonavis*) and one troodontid (*Jinfengopteryx*) among the vane-feathered paravians sampled have wing loading estimates at or below the 2.5 gcm^−2^ threshold for modern flapping flyers (Figs. 2a, S7, S8). Direct calculation of wing loading in *Mahakala* using an ultra-conservative wing area (Fig. 2b) shows that it fails the threshold of powered flight potential. The basal positions of the larger microraptorines *Tianyuraptor* and *Zhenyuanlong* imply a decrease in body size and an increase in relative forelimb length across Microraptorinae. This was confirmed quantitatively using parsimony-based ancestral state reconstruction of body mass (from femoral length) and forelimb length (from (humerus + ulna length)/femur length). This trend contrasts with multiple trends of body size and forelimb change (decreases as well as increases) in other dromaeosaurid lineages^2,71^. See Methods, Figs. 2 and S11-14. Using ancestral morphologies calculated with ancestral state reconstruction analysis, all fossil avialans sampled met the thresholds for powered flight seen in modern birds, suggesting that they had at least the potential for powered flight: wing loading values at or below 2.5 gcm^−2^ and specific lift estimates that exceed 9.8 Nkg^−1^. Direct calculations of wing loading and specific lift in our fossil avialan taxa show that only half of our minimum specific lift estimates (Po,m of 225 Wkg^−1^) failed the specific lift threshold for powered flight potential (Fig. 2b). The potential for powered flight we found in anchiornithids (*contra* ^18^) is supported by a reduced capacity for terrestrial running and greater emphasis on wing-based locomotion implied by the more proximal attachment of tail musculature, elongation of the acromion process, and more slender distal tibia found at the node shared between anchiornithids and traditional avialans^72^. The exception among anchiornithids is *Xiaotingia* which almost reached these thresholds for powered flight potential (Fig. 2). The potential for powered flight in anchiornithids should be treated cautiously though. This is because aspects of their anatomy may have affected their flight-relevant forelimb capabilities negatively e.g. lack of functionally asymmetrical feathers (char. 1.0); relatively short ulnae and humerii compared to derived avialans (char. 233.0; char. 262.2); limited pectoral musculature indicated by a weakly developed deltopectoral crest (char. 138.2) with an apex located closer to its proximal end (char. 684.2) and the lack of a bony sternum (at least in *Anchiornis*^56,73^). Paradoxically, *Xiaotingia* has a bowed rather than straight ulna, a feature linked with better takeoff potential in modern birds^74^. It also has a narrower radius (char. 438.1) like the aerodynamically capable *Microraptor* suggesting some potential benefits related to flapping-based locomotion. Alternatively, these features might yield mechanical advantages in contexts other than powered flight that deserve further investigation. The more active muscle-based shoulder stabilisation expected in basal birds is an anatomical limitation of powered flight potential that also needs to be considered because it would have used energy that could have otherwise been used for lift generation (their acrocoracoid process and/or its homologues is at or below the level of the glenoid: char. 342.0 of ^75^). A more passive and efficient intermediate condition of shoulder stabilisation did not appear until at least the node uniting *Jeholornis* and more derived birds, although the earliest members of this clade still lacked a bony sternum and modern arm flapping capabilities^72^ (the acrocoracoid process became elevated above the glenoid: char. 342.1; the strut-like coracoid appeared: char. 134.3; a more passive ligament system enables compressive forces to be transmitted from the wing to the sternum^75^). The stabilising role of a bony sternum is a synapomorphy of the more inclusive clade of *Jixiangornis* and more modern birds (char. 126.1). Shoulder stabilisation becomes even more passive and efficient at the node uniting *Hongshanornis* and more modern birds, when the humeral head becomes enlarged through the development of a proximal convexity (char. 352.1). However, modern-style arm flapping capabilities did not appear until later ornithurans^72^. Thus, fossil paravians that we suggest have the potential for powered flight likely did so less efficiently and with greater energy costs compared to modern birds.
2. *Aerodynamic force production requirements for flight* At the nodes Pennaraptora and Paraves, our wing loading estimates decrease and, to a lesser extent, our specific lift estimates increase (Fig. 2a). This coincides with a notable reduction in body size^3,71,72,76,77^, the appearance of pennaceous feathers^35,72,78^ (symmetrical at Pennaraptora^79^ and char. 456.1; asymmetrical at Paraves^72,79^) and a respiratory system more suited to higher intensity aerobic activity (advanced costosternal ventilator pump appearing among pennaraptorans^72^). Taken together, these findings support the suggested arm flapping capabilities of pennaraptorans^72^ as well as the potential for wing-assisted locomotion among paravians^72^. In other words, the data suggest that the wings of these taxa may have been used to assist in locomotion, such as running speed, turning, braking and jumping^18^. However, it is only at the node Paraves that either of our ranges of wing loading and specific lift estimates approach the minimal thresholds of powered flight potential (cf. initial aerial locomotion^72^) and only in Avialae, and a few independent lineages within Paraves (Unenlagiinae and Microraptorinae), where both thresholds are reached, thus indicating high probabilities of powered flight potential (Fig. 2). This supports the disconnect between the origins of pennaceous feathers and their incorporation into a flight-capable regime in non-avialan theropods^18,35^. Pennaceous feathers (char. 456.1) appear at Pennaraptora^79^ (but see ^47^) becoming asymmetrical in Paraves^72,79^. Asymmetrical forelimb feathers are found in *Microraptor* and are widespread among birds more derived than Anchiornithidae [char. 1.1]). This, in turn, lends credence to the hypothesis that pennaceous feathers and wings first evolved for non-flight purposes e.g. other wing-assisted locomotion^18^, display or egg brooding^80,81^.
3. *Physiological requirements for flight* Accounting for these constraints, among non-avialan paravians, only *Microraptor* and *Rahonavis* have specific lift estimates more than 9.8 Nkg^−1^, with no other non-avialan taxa having possibly been volant (Fig. 2a). Direct calculation of specific lift in *Microraptor* (Fig. 2b) shows that it passes the specific lift threshold for powered flight potential with our maximum estimate of specific lift using a Po,m of 287 Wkg^−1^. A wide range of plausible body mass estimates for *Microraptor* and *Rahonavis* derived from fossil measurements and 3D volumetric methods recovered the potential for powered flight: permutations including body masses up to double what would be expected for a living bird of similar span still retrieve a powered flight potential. For example, using the regression equations for wingspan vs mass calculated by Witton^82^ using a larger dataset of extant birds (n=96) or bats (n=102), we generate mass estimates of 0.445 and 0.323 kg respectively. Also, if we compare our estimate to a commonly used analog, the Common raven *Corvus corvax*, we find that similar-sized individuals have wing spans well in excess of one meter (data from: ^55,83^). Larger or more basal relatives of *Microraptor* and *Rahonavis* such as *Mahakala, Buitreraptor, Changyuraptor, Sinornithosaurus* and *Bambiraptor*, as well as all troodontids, show lower values of lift (Figs. 2, S9, S10). The less reliable calculation of ancestral wing loading and specific lift based on directly using those values from terminal taxa produced a similar picture (Fig. 2b). A wide range of deinonychosaurs showed wing loading values below the 2.5 gcm^−2^ threshold (Fig. 2). What is of particular interest is the subset of taxa below the wing loading threshold, but near or above the specific lift threshold (Fig. 2). Ancestral paravians shared several traits that presumably benefited the development of flapping-based locomotion, including smaller body size (^71,72^; Fig. 2), asymmetrically vaned feathers^72,79^, elongated and robust forelimbs^4,72^, a laterally orientated glenoid fossa articulation surface (char. 136.1; at equivalent node in ^2^: char. 138.1; related to the extension of the glenoid floor onto the external surface of the scapula: char. 138.1 in ^66^, char. 216.1 in ^29^, char. 122.1 in ^34^, char. 133.1 in ^35^ for node (*Jinfengopteryx* + Paraves)), a symmetrical furcula (char. 469.1), a laterally everted anterior edge of the acromion margin (char. 131.1) and elaborated brain regions associated with vision^72^. Although the origin of powered flight has been proposed at Paraves^72^, our data does not support this, but suggests the possibility of powered flight originating independently *outside* avialans. The unenlagiine *Rahonavis* is our strongest deinonychosaur candidate for flight potential, passing all wing loading and specific lift requirements (Fig. 2). This is consistent with the extremely elongated forearms of *Rahonavis* (ulna is longer than the femur as well as the tibia: Table 1 of ^84^) suggesting very large wings. The microraptorine *Microraptor* is another strong non-avialan candidate for flight, being below 9.8 Nkg^−1^ only for the strictest calculations (and even then displaying values approaching cut-off; Fig. 2). Its robust, asymmetrically feathered forelimbs controlled by muscles attached to a fused, ossified sternum^6^ support this. The vane asymmetry of its feathers though are less than the 3:1 vane ratio required for aeroelastic stability^61^, which might have limited *Microraptor* to relatively short flights (aeroelastic stability requires at least a 3:1 vane ratio). However, this is the case for many early avian taxa that otherwise seem flight capable. This is complementary to reconstructed aerodynamic prowess by several independent studies using traditional functional morphology^85^; physical^20-22,86^; and theoretical modelling^19,20^. Dececchi *et al*. ^18^ modelled launching in *Microraptor* (and other paravian taxa) similar to living birds. To further evaluate *Microraptor*’s candidacy for powered flight, we modelled its thrust-assisted launch potential under the alternative approach of ^87^, which used wing-generated thrust to supplement running takeoff in *Archaeopteryx.* We used their original parameters and calculated permutations that incorporated a larger flap angle and considered the effects of drag with both our model of *Microraptor* and the models of existing published studies^19,20,22^. In all cases, we found that *Microraptor* was capable of generating sufficient speed and flight forces for a ground-based takeoff and were within the range of values estimated for an arboreal launch^19,20,22^. Modelling approaches suggest that the 10% flight muscle ratio is probably underestimated for microraptorines (and *Archaeopteryx*^88^)^89^, although this low ratio is found in some living volant birds, such as particular owl species^90^. If we increase this ratio slightly to 13-15%, which is well within the range of living flying birds and is supported by volumetric modelling of these taxa, values well in excess of 9.8 Nkg^−1^ are achieved. For comparison, the average flight muscle fraction for living volant birds is around 18.4%, and this average is somewhat elevated by highly derived forms such as passerines that often have high flight muscle fractions^90^. Such promising flight potential provides a compelling context for interpreting unusual, potentially flight-related anatomies in more detail (e.g. the elliptical fenestra of the deltopectoral crest found in *Microraptor* and the volant early birds *Confuciusornis* and *Sapeornis*^16,91,92^). See Supplementary Note 3 for additional information about microraptorines. Although other paravian taxa such as the troodontid *Jinfengopteryx* and the dromaeosaurids *Bambiraptor, Buitreraptor, Changyuraptor* and *Mahakala* are also close to these thresholds, they never surpass them despite the generous wing and flight muscle ratio reconstructions adopted (Fig. 2). Even though they did not pass the specific lift threshold, their high scores as well as low wing loading values makes these taxa – particularly the microraptorine *Changyuraptor* – deserving of further study from the flight potential perspective, using more fine-grained techniques and modelling. This will distil the extant nature of the changes that are necessary to transition from non-volant flapping based locomotion to active flight. The recent suggestion of a short-armed clade at the base of Dromaeosauridae^93^ supports the idea that flight capability is not ancestral to paravians.

Phylogenetic distributions of wing loading and specific lift (see Figs. S7-10 and Methods) combined with osteological, integumentary and body size changes provides a more holistic and integrate view of the origin of powered flight. The robust phylogenetic context allows us to examine the evolutionary transitions of powered flight requirements from the perspective of anatomy, aerodynamic force production and muscle physiology. Of these three categories of flight requirements, we are most confident that some small non-avialan paravians had the required anatomical competency for flight. We are highly confident that aerodynamic force production was sufficient for flight in *Microraptor, Rahonavis*, and early birds. Because muscle physiology is not known for fossil taxa, and because our specific lift estimations must necessarily make more assumptions than the other aspects of the analysis, we are less confident regarding the precise patterns of specific lift evolution recovered in the analysis. However, our results do show that, for conservative muscle power outputs, some of the non-avialan paravian taxa could likely fly, even if only briefly at lower power outputs. All permutations for *Rahonavis* suggest powered flight potential as do 9 of our 12 permutations of the *Microraptor gui* model using 10% muscle mass and all 12 using 13% muscle mass as estimated by Allen et al. ^94^. From these results, we suggest that muscle physiology might have been the limiting constraint for flight in early paravians. Under this model, any time muscle physiology crossed the critical power output boundary, flight could have originated – and this could have happened multiple times. Our results allow us to test a number of other hypotheses relating to 4 areas:

1. *Muscle size and physiology:* We reject the hypothesis that flight muscle fractions above 10% would be required for large-winged non-avian pennaraptorans to engage in powered flight. We further reject the hypothesis that flight muscle physiologies outside of those seen in modern birds would be required for large-winged non-avian pennaraptorans to engage in powered flight.
2. *Wing area:* We reject the hypothesis that large-winged non-avialan pennaraptorans would have been prevented from flight on account of insufficient wing area relative to body mass. Only under the most extreme body mass estimates for large-winged non-avialan pennaraptorans do we retrieve wing loading results above the powered flight thresholds observed in living birds. For example, even using the heaviest mass estimate per Allen *et al*. ^94^ of 1.59 kg for *Microraptor gui* and the lowest wing area estimate of Chatterjee and Templin^19^, based on the incomplete estimate of feather length, we still obtain wing loading values of 1.69 gcm^−2^, well below the 2.5 gcm^−2^ maximum and similar to values seen in adult chukar partridges^83^ and turkeys^95^.
3. *Duration of aerial behaviours:* We cannot reject the hypothesis that powered aerial behaviours in large-winged non-avialan pennaraptorans were typically brief in duration. While the gross wing structure in large-winged non-avian pennaraptorans appears to be very similar to that of living birds, the structure of individual feathers suggests that aeroelastic instability in early taxa may have reduced wing performance. In some permutations of our estimated parameter set, recovery of power flight potential in large-winged non-avialan pennaraptorans was dependent on a portion of the flight muscle mass being anaerobic. For example, all anchiornithids at a 10% muscle fraction require muscle outputs of minimally 250Wkg^−1^ to achieve sufficient lift for takeoff (except *Xiaotingia*, which never achieves sufficient lift). In these cases, low muscle endurance would necessitate short-ranged aerial behaviours.
4. *Powered flight potential across Paraves:* We cannot reject the hypothesis that large-winged non-avialan pennaraptorans potentially had powered flight, and that of some kind of potential evolved multiple times among paravians. The majority of our parameter permutations recover some level of powered flight potential in large-winged non-avialan paravians. Based upon these results, we feel that it is likely that powered flight evolved multiple times from a range of paravian ancestors that were already nearing powered flight potential: twice in Dromaeosauridae, once or twice in Avialae [depending on character optimisations for *Xiaotingia*], and potentially once in Troodontidae [if more capable examples than *Jinfengopteryx* are found] (Fig. 2)), but originated neither with Avialae nor at a single node within Paraves. We consider this more likely than a scenario in which the body densities or muscle physiologies of non-avialan pennaraptorans were far outside those measured in living birds. These potential powered flyers are all associated with size reductions as well as forelimb elongation (in this study and in ^18,71^). Notable anatomical differences between subclades (e.g. the absence of ossified sterna in troodontids and their presence in dromaeosaurids^73,96^) suggest these independent origins of flight are not entirely parallel – i.e. they do not share the same anatomical starting points and may have achieved functional flight in different ways. This also appears to be the case for non-paravian pennaraptorans, as suggested by the bizarre membranous wings of the scansoriopterygid *Yi qi*^34^. The relative paucity of preserved troodontid forelimbs compared to those of dromaeosaurids hinders some of these reconstructions, but known forelimb differences have intriguing implications for the evolution and ecology of paravian powered flight.

**Fig. 2.**
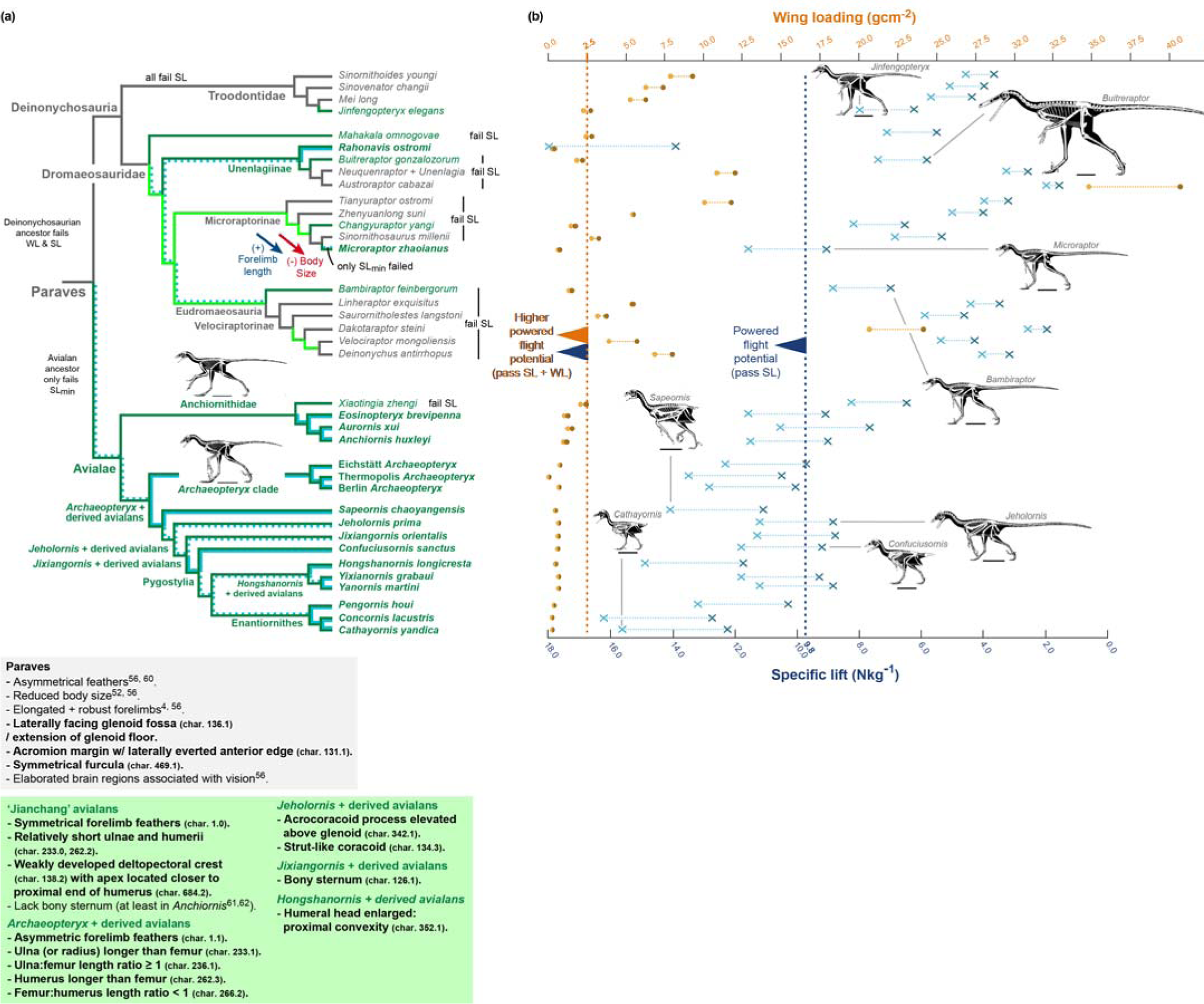
Maximum and minimum estimates of wing loading and specific lift across Paraves and flight-relevant synapomorphies. (a) *Parsimony-based ancestral state reconstruction analysis of paravian wing loading and specific lift using the new phylogeny.* Powered flight potential in an ancestor or terminal taxon as suggested by wing loading estimates above the 2.5 gcm^−2^ threshold (Figs. S7, S8): marked in dark green shading if relating to the maximum reconstructed ancestral value (i.e. minimum powered flight potential; minimum wing area and maximum body mass); marked in both dark green and light green shading if relating to the minimum reconstructed ancestral value (i.e. maximum powered flight potential; maximum wing area and minimum body mass). Grey shading denotes wing loading estimates below the 2.5 gcm^−2^ threshold. Powered flight potential in an ancestor or terminal taxon as suggested by specific lift values above the 9.8 Nkg^−1^ threshold (Figs. S9, S10): marked in stippled light blue shading if relating to a maximum estimate (SL_max_ uses maximum muscle mass-specific power output (Po,m) of 287 Wkg^−1^); marked in light blue shading if relating to both the minimum and maximum estimates (SL_max_ and SL_min_ uses Po,m of 287 Wkg^−1^ and 225 Wkg^−1^ respectively). Specific lift estimates below the 9.8 Nkg^−1^ threshold are not marked. The figure includes the trends of increased forelimb length (dark blue arrow) and decreased body size (red arrow) along the microraptorine lineage. A list of flight-relevant synapomorphies is provided in the text boxes (synapomorphies in bold font are from this study, whereas unbolded ones are from the literature). See Methods for additional information. Skeletal reconstructions used with the permission of Scott A. Hartman (scale bar = 10cm). (b) *Maximum and minimum estimates of wing loading and specific lift for each terminal taxon.* Maximum and minimum estimates of wing loading calculated using conservative and ultra-conservative wing areas (light orange and brown dots respectively). Maximum and minimum estimates of specific lift calculated using a broad range of Po,m values (287 Wkg^−1^ and 225 Wkg^−1^). These terminal taxon values were not calculated from ancestral morphologies using ancestral state reconstruction analysis. See plotted values and their derivation in Supplementary Files at www.palaeopittman.com/flightoriginspaper [password for review process: flightorigins]. Skeletal reconstructions used with the permission of Scott A. Hartman (scale bar = 10cm).

Our analysis suggested multiple origins of powered flight from differing initial conditions with some members exhibiting some capacity for wing-based locomotory assistance that, although not flight capable, may have assisted non-volant behaviours^18^. This implies that Paraves, in general, may have been experimenting with wing-assisted, non-volant behaviours to expand into locomotory repertoires otherwise unexplored by Late Jurassic and Early Cretaceous vertebrates. These include high-speed running and starts, leaping, rapid braking and turning, and dynamic balance. Emphasis on some of these behaviours in different paravian, and even pennaraptoran, clades may have presented opportunities for diverse ecological niches for these agile taxa. Only when some clades evolved smaller body sizes did these independent biomechanical repertoires, adapted for high speed terrestrial or scansorial locomotion, became capable of powered flight. This evolutionary scenario emphasises further examination of the non-volant, large-bodied paravians with the goal of estimating their differing anatomical and biomechanical specialisations. The results presented here suggest paravians were exploring a wider range of locomotory niches than previously appreciated and may have set the stage for the origin of birds and powered flight from a rapidly evolving, highly diverse suite of locomotory and ecological experimentations.

## Methods

### Phylogenetic dataset

The coelurosaurian theropod dataset of Brusatte *et al.* ^1^ (the most recent version of the Theropod Working Group dataset [TWiG dataset]) was significantly expanded with data pertinent to paravian phylogeny, especially data concerning dromaeosaurids. Nine dromaeosaurid terminals were added to this expanded version of the TWiG dataset for the first time, including the Late Cretaceous microraptorine IVPP V22530, *Changyuraptor, Zhenyuanlong, Luanchuanraptor, Acheroraptor, Linheraptor, Yurgovuchia, Dakotaraptor* and *Velociraptor osmolskae*. The current dataset has thirty-one dromaeosaurid taxa, including all valid genera that have been included in previous phylogenetic analyses, except for *Pyroraptor* (represented by a fragmentary specimen lacking recognisable synapomorphies^2^). Codings of many other dromaeosaurids, troodontids and basal avialans have been revised, including *Anchiornis, Aurornis, Eosinopteryx* and individual specimens of *Archaeopteryx*. Codings for several non-paravian maniraptorans have also been revised or added for the first time e.g. the scansoriopteryid *Yi*. See Supplementary Note 4 for a complete listing of all coding changes. All taxa were coded based on first-hand examinations, relevant literature and photographs. Some codings for the newly included taxa e.g. *Acheroraptor, Yurgovuchia, Dakotaraptor* and *Yi* were also adopted from non-TWiG datasets^29,31,33,34^. Four *Archaeopteryx* specimens (Eichstätt, Berlin, Haarlem and Munich) were re-examined first-hand using Laser-Stimulated Fluorescence (LSF) imaging^97^, revealing additional anatomical details.

The character list of Brusatte *et al*. ^1^ consists of 853 characters compiled from multiple sources. A new character state was added to character 229 (to reflect a potential synapomorphy of *Archaeopteryx* specimens on the ischium) and character 744 (to reflect the variations of pedal phalanx II-2 in deinonychosaurians) (see Supplementary Note 5).

### Automated phylogenetic analysis

The phylogenetic analysis was carried out with TNT version 1.5^98,99^. In order to make the analysis fully reproducible, a master script was used to automate thorough searches, as well as the subsequent diagnosis and characterisation of results. All the scripts and batch files for initial analysis, diagnosis, and other tasks, are available (with full descriptions) in the Supplementary Methods.

### Reconstruction of wing loading and specific lift values

Two criteria – wing loading and specific lift – were taken from theoretical and *in vivo* work on extant avialans and applied to fossils; they present easily testable benchmarks to discern volant from flightless taxa^44-46,68^. For taxa without preserved complete primary feathers, feather length was modelled on closely related taxa and wing area was calculated based on the methods presented in Dececchi *et al*. ^18^ (See spreadsheet in Supplementary Files at www.palaeopittman.com/flightoriginspaper [password for review process: flightorigins]). Wing span was taken as 2.1 times the summation of the lengths of the humerus, ulna and metacarpal II and the longest distal primary^18^. Wing chord was taken as 55% of the longest distal primary length, a modification of the methodology used in^18^. This is because it better reflects the differences between primary and secondary lengths seen in *Microraptor*^100^ and produces wing area estimates that are within less than 1% of those measured by Yalden^101^ for *Archaeopteryx* and by Lü and Brusatte for *Zhenyuanlong*. To improve optimisation of the data we screened these coelurosaurs from 77 to 43 taxa based on their presence of vaned feathers which are integral to the production of aerodynamic forces; terminals for which feather condition is unknown were considered to have the same state as their ancestor, which is the condition predicted by our phylogenetic hypothesis (if absent marked as ‘---’). The same mapping method for reconstructing dromaeosaurid body size and forelimb (section directly above) was used to reconstruct wing loading and specific lift values.

#### Wing loading

Meunier and others demonstrated that volant extant birds always have wing loading values below 2.5 gcm^−2 44,69,102-105^, and so the present study deems a fossil taxon with values above 2.5 gcm^−2^ as certainly flightless. Fossil taxa with values above 2.5 gcm^−2^ are seen to have had the potential for powered flight. Wing loading is based on body mass estimated as per above (kg; see ‘Trends in body mass change and forelimb length’ in Methods) over wing area (cm^2^).

#### Specific lift

In the case of specific lift, the cut-off used to identify fossil taxa with the potential for powered flight is 9.8 Nkg^−1^ (gravity), as used by studies involving volant extant birds^45,46^. In practice the value is slightly *greater* than 9.8 Nkg^−1^ since some lift is oriented as thrust in powered flyers^87^. Specific lift is based on Marden’s model^45^:

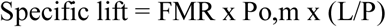

Where FMR is the flight muscle ratio which was assigned at a constant value of 10% of body weight across all taxa examined here. This is at the lower range of the values seen in volant birds and is likely a significant overestimation for all non-paravian taxa, though lower than those for *Archaeopteryx* and *Microraptor* based on recent 3D modelling work^89^. Po,m is the maximum muscle mass-specific power output based on values from extant birds. As Po,m is unknown for non-avialan theropods, two separate calculations were made that bracket the range of Po,m values that could have reasonably been expected (225 and 287 Wkg^−1^; see Figs. S9 and S10 as well as TNT scripts and script results in Supplementary Files at www.palaeopittman.com/flightoriginspaper [password for review process: flightorigins]) to reconstruct minimum and maximum powered flight potential. L/P (lift/power) is calculated from:

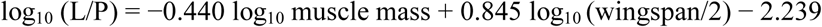

### Uncertainty quantification and estimation confidence

One potential weakness of our modelling approach is the sensitivity to scaling assumptions in the assessment of locomotor performance. This sensitivity does not affect taxa recovered as far below thresholds for volancy, but it could potentially affect conclusions for those taxa recovered as performing near thresholds for volant behaviour i.e. near powered flight potential. To address this, we used an iterative resampling method in which we varied the starting parameters and reran the analyses for taxa recovered as having performance estimates near threshold values. We found that our model was most sensitive to assumptions regarding specific lift, and so we focused resampling on varying FMR. As noted above, Po,m was automatically varied for all taxa by performing calculations at two values that encompass the range of maximum power outputs measured by prior teams (see section above ‘Reconstruction of wing loading and specific lift values’. Error in mass estimation was found to be less critical, with marginal performance taxa typically requiring both a significant deviation in wing area and a significant deviation in body mass from the expected values to change our expectations of volancy. However, varying body mass by applying standard errors from the mean as scalars is arguably not the most robust method, since this ignores the underlying frequency distribution. To further validate masses for the most critical taxa (particularly *Microraptor*), we validated our estimates against wholly independent methods of mass estimation, including those derived through 3-D computer and displacement methods (see ‘Trends in body mass change and forelimb length’ section above). Validating our mass estimates against volumetric-based estimates is a particularly robust option because it allows us to eliminate extraneous potential minima and maxima that would result in unrealistic body densities (i.e. those well above or below those measured for living birds).

### Trends in body mass change and forelimb length

Paravian body masses were calculated from femoral length measurements using the empirical equation of Christiansen & Fariña^106^:

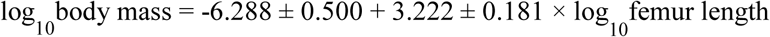

This methodology is a widely used estimator for body size across Theropoda^71^. While limb bone circumference has been shown to be a more accurate proxy of theropod body size^3,107^, this measurement was not available in many important Chinese paravian taxa because their long bones are crushed or flattened on mudrock slabs (a survey of ∼1000 specimens covering dozen of species failed to recover reliable circumferences). Thus, the femoral length proxy was adopted because the measurement itself is available across our sample and because it has been widely used in previous theropod literature. As *Microraptor* is critical for our analysis we compared our mass value to one generated from an estimate of femoral circumference using the empirical equation of Campione *et al*. ^108^ as well as comparisons to mass estimates generated through 3-D computer and displacement methods^19,22,89^. All of these produce similar estimates to the one obtained using Christiansen & Fariña^106^ (See Supplementary Tables 1, 2). Among all possible reconstructions, the maximum possible increases in size (or the minimum decrease) were calculated. See Supplementary Methods for mapping methodology.

### Data availability

The data that support the findings of this study are available from the corresponding author upon reasonable request. The data reported in this paper are detailed in the main text and Supplementary Information.

## Acknowledgments

This study was supported by the Research Grant Council of Hong Kong’s General Research Fund (17103315 to M.P.). It was also supported by The University of Hong Kong (Faculty of Science (to M.P.), University Research Committee Postdoctoral Fellow Scheme (to M.P. for R.P.), Conference Support (to M.P.) and Seed Fund for Basic Research (two awards to M.P.)), the National Science Foundation of China (41688103, 41120124002 and 91514302 to X.X.), and a Proyecto de Unidad Ejecutora from CONICET (PUE0070 to P.A.G). This study benefitted from discussions at the International Pennaraptoran Symposium, held at the University of Hong Kong and supported by Kenneth HC Fung and First Initiative Foundation. Scott A. Hartman is thanked for the use of his skeletal reconstructions in figures 1 and 2.

## Author contributions

M.P., P.A.G., X.X. designed the project with input from T.A.D. and R.P.. All authors performed the research, wrote the manuscript and gave feedback on the manuscript.

## Additional Information

**Supplementary Information** accompanies this paper

